# Family aggregation analysis reveals a heritable background of equine grass sickness (dysautonomia)

**DOI:** 10.1101/592253

**Authors:** Boglárka Vincze, Márta Varga, András Gáspárdy, Orsolya Kutasi, Petra Zenke, Ottó Szenci, Ferenc Baska, Alan Bartels, Sándor Spisák, Norbert Solymosi

**Affiliations:** Laboratory of Genetics, Unit of Animal Breeding and Genetics, Department of Animal Breeding, Nutrition and Laboratory Animal Science, University of Veterinary Medicine, 1078-Budapest, Hungary; MTA-SZIE Large Animal Clinical Research Group, Hungarian Academy of Sciences, 2225-Üllő, Hungary; Department and Clinic of Zoo and Wildlife Medicine, University of Veterinary Medicine, 1078-Budapest, Hungary; Department of Medical Oncology, Dana-Farber Cancer Institute, Harvard University, 02215 Boston, Massachusetts, USA; Centre for Bioinformatics, University of Veterinary Medicine, 1078-Budapest, Hungary

**Keywords:** grass sickness_1_, dysautonomia_2_, heritable_3_, equine_4_, horse_5_

## Abstract

Equine grass sickness (also known as dysautonomia) is a life-threatening polyneuropathic disease affecting horses with approx. 80% mortality. Since it’s first description over a hundred years ago, several factors including phenotypic, environmental, management, climate, and intestinal microbiome) have been associated with increased risk of dysautonomia. But despite the extensive research on dysautonomia, it’s causative factors have yet been identified. A retrospective pedigree and phenotype based genetic epidemiological study was performed to analyze the associations of disease occurrence and the kinship in a Hungarian large scale stud. The pedigree data set containing 1233 horses with 49 affected animals was used in the analysis. The first finding was that among the descendants of some stallions the proportion of affected animals are unexpectedly high, with a maximum of 25% of a stallions descendants affected. Animals with affected siblings have higher odds to be a case (OR: 1.27, 95% CI: 1.01-1.57, p=0.033). Among males in the affected population the odds of dysautonomia is higher than in females (OR: 1.76, 95% CI: 0.95-3.29, p=0.057). Significant familial clustering was observed among the affected animals (GIF p=0.001). Further subgroups were identified with significant (p<0.001) aggregation among close relatives using kinship-based methods. Our analysis of the data and the observed higher disease frequency in males suggests that dysautonomia may have X-linked recessive inheritance as a causal factor. This is the first study providing ancestry data and suggesting a genetic contribution to the likely multifactorial causes of the disease.

## 1 Introduction

Equine dysautonomia (also known as EGS/equine grass sickness) is a life-threatening disease, with an unknown etiology, affecting horses. Approximately 80% of the affected animals die within days after recognition of the first clinical signs; the mortality is reported as 100% in acute and subacute form and 50% in the chronic form (McGorum and Pirie, 2018). The acute form of the disease leads to death in 100% of cases. Horses with chronic features of the disease have a shortened survival, and a 50% mortality. This polyneuropathy affects both the central and peripheral nervous system and leads to severe clinical symptoms, as well as gross and microscopic pathologic alterations in the ganglions and neurocytes localized primarily in the gastrointestinal tract. Since its first recognition in 1909 (Tocher et al., 1923), multiple cases have been reported around the world (McGorum and Pirie, 2018).

Despite significant effort and research, neither the causative reason, nor the pathomechanism of the disease are well-understood. The most widely accepted hypothesis is a possible toxicoinfection of *Clostridium botulinum* type C, which was detected in the dissected horses’ gut microflora (Hunter et al., 1999). However, studies performed to vaccinate horses against the bacteria have yielded mixed results, as successful (McCarthy et al., 2004) and unsuccessful (Nunn et al., 2007a) vaccination attempts have been reported Decades of research have associated several factors with an increased or decreased risk of the disease (Newton et al., 2004, McCarthy et al, 2004, Wood et al., 1998, Doxey et al., 1991a, 1991b, Edwards et al., 2010; Wylie et al, 2014, 2016). Among these factors: phenotypic (age, body condition), environmental (soil nitrogen content, grass, number of horses, presence of birds, ruminants, younger animals, etc.), management (grazing, movement, stress, anthelmintic use, supplementary feeding, etc.) and climate (cool, dry weather, frosts, increased sun hours, maximum temperature, snowing) (McGorum and Pirie, 2018). The disease has different rates of prevalence between countries, but EGS can cause high morbidity (10-25%) and high mortality (50-100%) rates within a single herd resulting in significant economic losses. Until recently, family patterns and pedigree analysis could not performed as the majority of infected animals were of different origin.

The aims of the present study were a.) to perform a pedigree analysis on an study population affected by EGS, b.) to detect signs of heritability of EGS, and c.) to provide evidence for genetic predisposition to EGS.

## 2 Materials and Methods

### 2.1. Study population

The original goal of the study was to investigate the problem of equine deaths in Hungary, and to reserve important genetic lines for horse breeding. Horses have been kept and bred here in Hungary for nearly 200 years without large historical problems. But every year, from 2001 onwards, 0-11 horses die due to unknown causes, resulting in significant economic and population-wide losses. Initial autopsy and pathological findings performed in 2001-2002 reported a diagnosis of EGS. Reports showed that before their death, all of the affected animals were in a normal body condition, without advance indication of the severe colic symptoms typical of EGS. Multiple attempts have been made to treat affected animals, but have failed at preventing losses.

Detailed description of the first findings can be found elsewhere (Vörös et al., 2003, Schwarz, et al., 2012).

The stud farm for this study is situated in Northeastern Hungary among the Bükk Mountains. The assessed population consists of 70 broodmares, 12 stallions, approx. 150 weanlings and yearlings from the last breeding seasons and 40 non-breeding horses. A total of approx. 280 horses are kept in 3 places close to each other. The affected groups are kept in separate facilities based on their sexes. The affected animals are among the 1-4 years old young horses from both sexes. Horses were kept and fed under the same conditions (approx. 2 kg oats and 6 kg grass hay daily with unlimited access to water). The mares and younger horses (age<4 years) were moved to a pasture (located approx. 3 km away) twice daily.

After the initial outbreaks of the disease in 2001, multiple attempts have been made to investigate the possible reason for the significant losses. Between 2001 and 2017, the water, all the feeds, the grass and grass hay, the soil, and other environmental conditions have been examined for toxins but none of them could be detected as a causative.

### 2.2. Data Collection

Pedigree data for the analyzed population was provided by the Hungarian Association of Lipizzaner Horse Breeders and the Directorate for Animal Breeding (by Barnabás Szabó, National Food Chain Safety Office). Health and pathologic data was collected from the stud records of the affected stud.

### 2.3. Statistical analyses

For testing the sex independence of the EGS condition, Fisher’s exact test was used (Agresti, 2007). The confidence interval of prevalence was estimated by Wilson method (Agresti and Coull, 1998). The family aggregation was analyzed by genealogical index analysis (Hill, 1980), kinship sum test, kinship group ratio test (Rainer et al., 2016; Dinya and Solymosi, 2016) and familial incidence rate test (Kerber, 1995). The method of Wellmann et al., (2012) was applied to estimate the time at risk required by the familial incidence rate test, as for numerous animals the amount of time spent in the stud was unknown. All statistical analyses and visualization were performed in R (R Core Team, 2018).

## 3 Results

Among the 1233 animals in the study population, 49 were affected with EGS. The number of registered cases ranged yearly between 1 to 11 since 2001(Figure 1). The mean age of diseased animals was 2.45 year (SD: 0.54). Comparing the rate of disease occurrence in males (28/539) and females (21/694), we found that the males have higher odds to be affected (OR: 1.76, 95% CI: 0.95-3.29, p=0.057). Analyzing the involvement of descendants of particular stallions the overall prevalence of the disease were 3.97% (95% CI: 2.99-5.22). Some males had larger affection frequency among his successors, for instance among the 32 descendants the stallion No. 719 (Figure 2) there was 7 affected (21.9%, 95% CI: 10.46-38.95).

**Figure 1.**
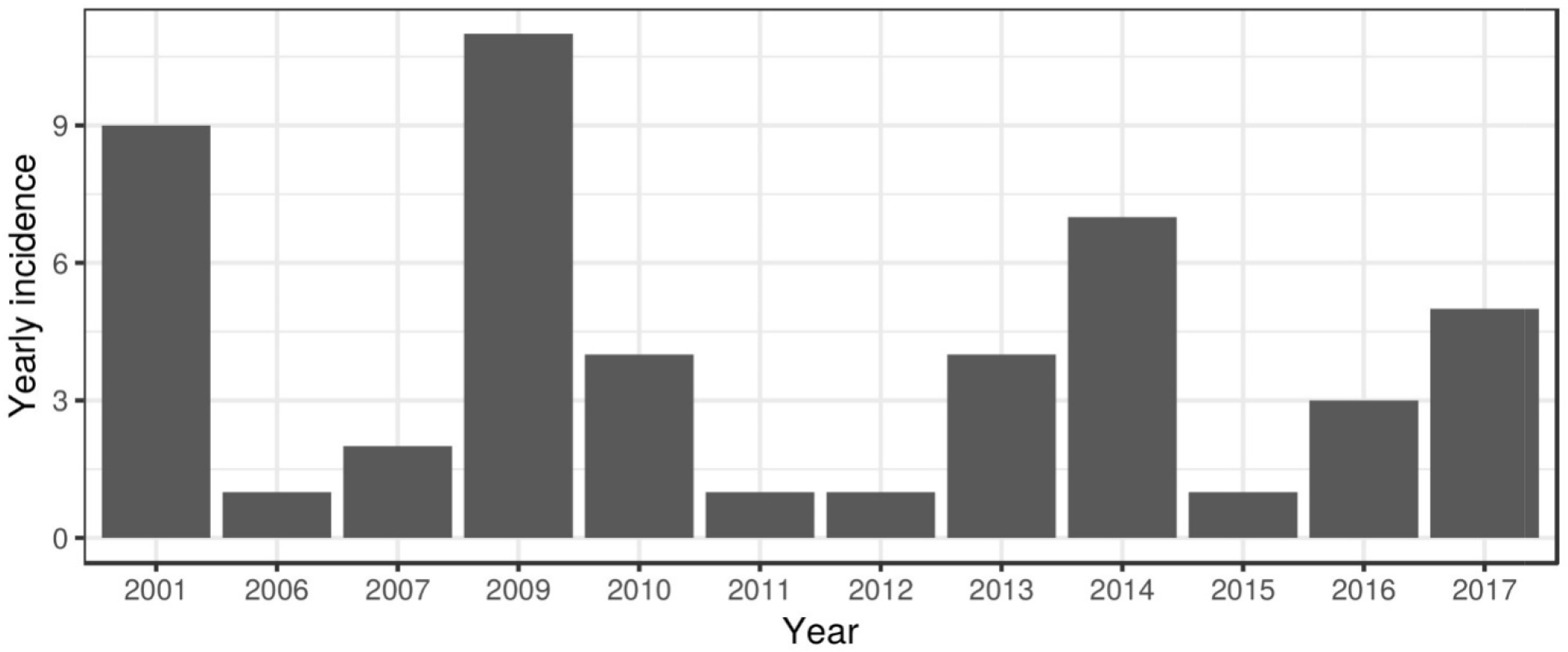
Distribution of the occurrence of diseased animals by year.

**Figure 2.**
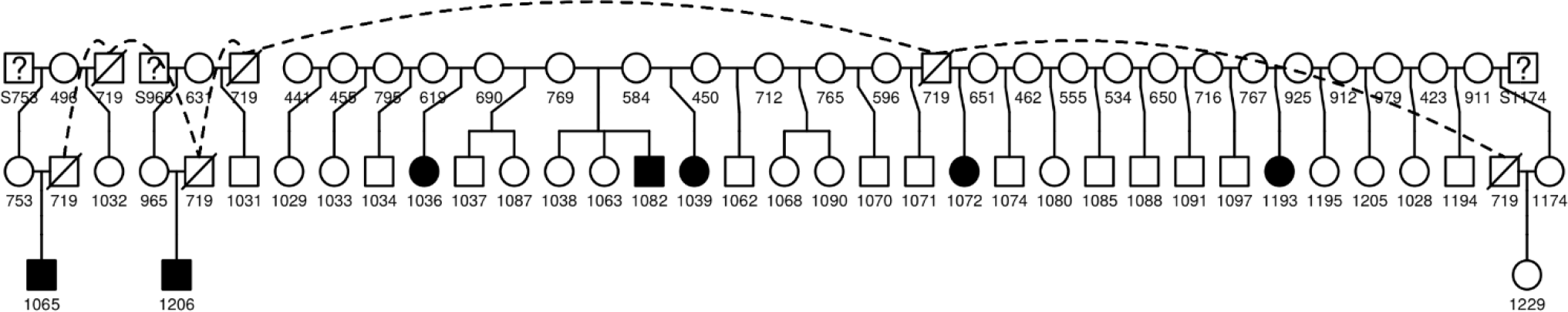
The family tree of stallion Nr. 719 (crossed box), the black members are affected, and the whites are not affected.

The mares’ successor was also analyzed, but there was no similar large outcome frequency found. With the genealogical index analysis we found that the mean kinship is significantly higher (GIF: 3130.71, mean kinship: 0.031, p=0.001) among the affected animals than it’s expected value (Figure 3).

**Figure 3.**
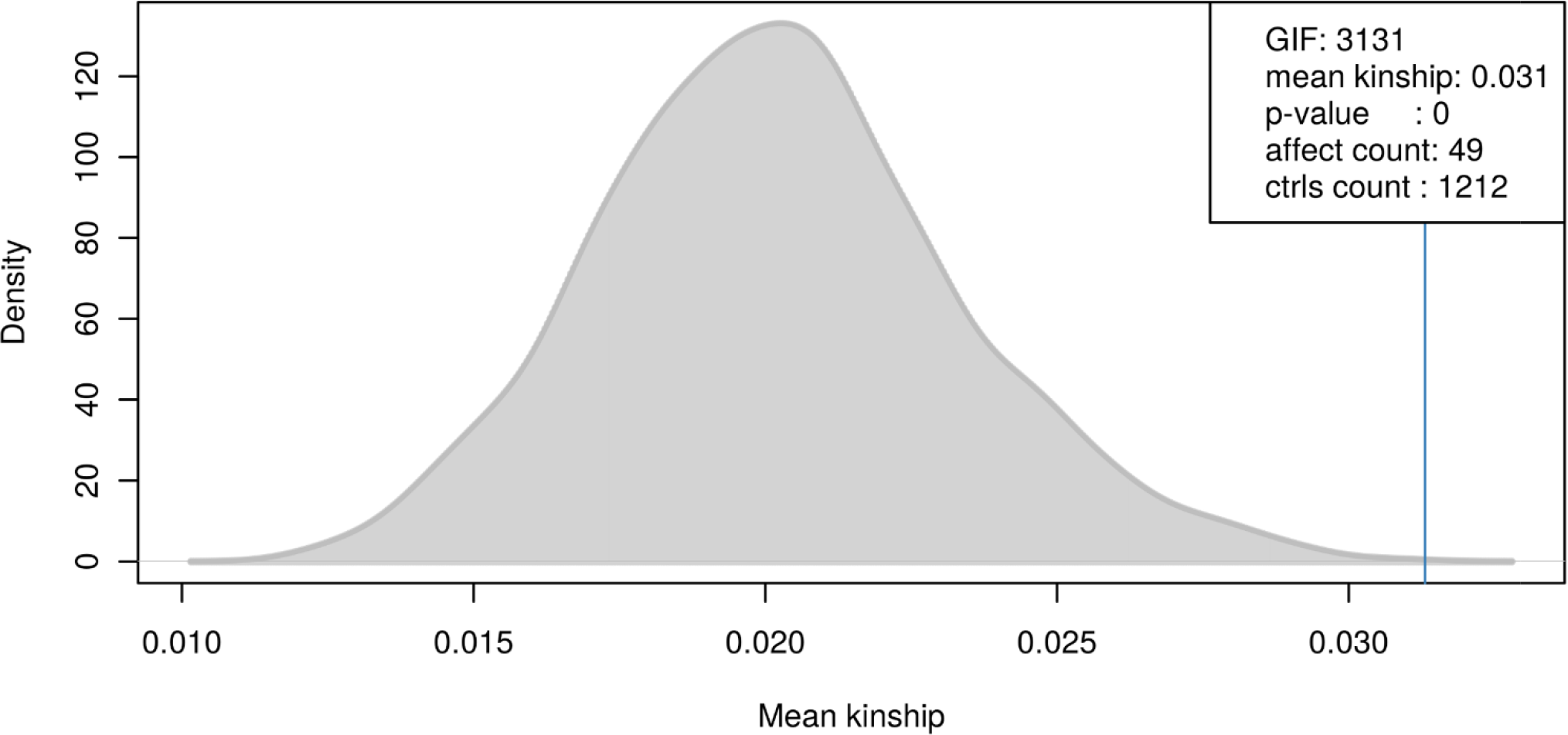
Results of genealogical index analysis: blue line represents the observed, the grey area represents the expected mean kinships among the affected animals.

By performing familial aggregation test based on the kinship sums, and selecting individuals using the adjusted p-values because of multiple testing, we found the Nr. 933 animal as the member of pedigree highly connected to other affected members. Checking the ascendants of this mare, her parents, grandparents and grand-grandparents had consanguineous matings. With the kinship group ratio test we found three subsets in the pedigree where the observed number of the affected individuals was significantly higher (p adj. < 0.001) than expected. By familial incidence rate test we found 82 individuals having significant (p adj. < 0.001) FIR-value, the highest FIR-value (0.019) belongs to the Nr. 719 animal.

## 4 Discussion

Equine grass sickness (or dysautonomia, EGS) is a neurological disease affecting the autonomous and peripheral nervous system in horses of unknown origin. Nowadays it occurs worldwide except Africa and Asia. In the years between 2000-2009, 141 cases have been reported annually on average in Great-Britain (Wylie et al., 2011).

The disease can be present in three clinical forms but with different frequencies (47% acute, 20% subacute and 33% chronic) according to McCarthy et al., (2001). The fight against grass sickness is important among equine clinical studies because of the very high mortality (50-100%) rate among affected animals.

The diagnosis of EGS is based on the clinical and histopathological findings occurring in the diseased animals and is well described elsewhere (Cottrell et al., 1999, Milne 1991, Pirie et al., 2014).

In the last century, many researchers attempted to characterize the etiology and pathophysiology of the disease, but even the infectious background could not be proven; nobody was able to provoke the symptoms by trials/experiments. Various predisposing factors have been analyzed (age, breed, stress, change in environmental conditions, season, weather, etc.) and numerous possible causative agents have been investigated (viruses, bacteria and their toxins, mycotoxins, toxic plants, chemicals, nutritional factors, insects, bugs, antiparasitic drugs, etc.) by researchers (McCarthy et al., 2001).

The most common theory among researchers regarding the cause of the disease is the ‘botulism-theory’. Many studies concluded that *Clostridium botulinum*, which produces toxins in the horses’ intestines, can cause the disease and some investigations showed higher prevalence of this bacteria in the affected horses’ samples (Wood et al., 1998 and 1999, Murray et al., 1994, Nunn et al., 2007a, 2007b) and that the lesions caused by this toxin (BoNT-C) were similar in the affected horses’ nerve-endings as described in *in vitro* rat tissue studies (Williamson et al., 1995). Systemic antibody levels after vaccination against *C. botulinum* showed favorable results in some case-control studies (Hunter et al., 1999, Hunter and Poxton, 2001). Between 2013-2016, a vaccination trial was performed in the study population. After a 3-element vaccination and check of the immunoglobulin-levels, 2 vaccinated foals died at age 3 years. Therefore the vaccination did not help to avoid the occurrence of EGS in this stud (personal communication – Dr. Orsolya Kutasi).

The disease is most prevalent in horses between 2-7 years old with a peak at the age of 4-5 years (McCarthy et al., 2001). In the present study, the mean age of the diseased animals was 2.45±0.54 years which agrees with other reported findings. Although there was no sex-predisposition in the previous studies (Wylie 2014), we found a higher odds (OR: 1.76) in males to be affected. This indicates that a possible X-linkage should be considered. The X-linked association could not be completed because of the high occurrence of consanguineous matings in the pedigree.

The overall disease prevalence was 3.97% in the study population which agrees with previous studies in Great-Britain (Cottrell, et al., 1999), but some stallions (e.g. Stallion Nr 719 had 7/32 [21.9%] diseased foals) had much higher number of affected descendants than the population average.

A familial aggregation test is the first step in the identification of genetic determinants of a disease (Matthews et al., 2008). If aggregation is found, more refined genetic studies are needed to identify a specific alteration in genetic material (e.g. mutation). In this study, one mare (Nr. 933) was found to be highly connected to other affected animals based on kinship analysis. Evaluating the pedigree of mare Nr. 933 we found numerous consanguineous matings leading to inbreeding. The familial incidence rate test showed a significant FIR (familial incidence rate)-value among affected animals; therefore we may conclude that inbreeding causes the aggregation (phenotypic appearance) of equine grass sickness.

To summarize, it is always a hard task to determine whether a disease is heritable. A thorough pedigree analysis should be performed as the first step to examine a possible aggregation in the family. The statistical analysis methods differ and none of them can verify a genetic background individually, therefore we used several methods during the investigation. Based on presented results, the authors think that the solution is likely difficult – the predisposition might be inherited and maybe an additional factor is needed for the clinical signs to occur. In the literature of equine grass sickness and in the study population, the stress (e.g. change in environment) as a factor is likely to be present before the clinical signs develop. Recent transportation, feed change, cool dry weather, irregular frosts (McGorum and Pirie 2018), storm are mentioned as a possible factor predisposing horses for EGS.

The question which now comes to mind is whether the disease or it’s predisposition can be identified by an alteration in DNA sequence. The aggregation among relatives as a consequence of inbreeding in the affected study population has been shown here. To improve our understanding of the genetic background of the disease a genome-wide association study (GWAS) should be performed to find differentiating genes, polymorphisms. If such causative features are found, further works may lead to a practically applicable genetic test. If such a test was available, the genetic predisposition will be identifiable in horses and a breeding selection based on test results can be done to avoid the phenotypic appearance of equine grass sickness in the near future.

## 5 Conflict of Interest

The authors declare that the research was conducted in the absence of any commercial or financial relationships that could be construed as a potential conflict of interest.

## 6 Author Contributions

Conceptualization: András Gáspárdy, Boglárka Vincze and Norbert Solymosi

Data curation: Boglárka Vincze, Márta Varga, Ferenc Baska, Orsolya Kutasi, Petra Zenke Investigation: Boglárka Vincze, Márta Varga, Norbert Solymosi

Methodology: Norbert Solymosi and Sándor Spisák

Project administration: Ottó Szenci, András Gáspárdy

Writing – original draft: Boglárka Vincze, Alan Bartels and Norbert Solymosi

## 7 Funding

Supported through the New National Excellence Program of the Ministry of Human Capacities Hungary (Ms. Márta Varga).

## 8 Acknowledgments

The Authors would like to thank Barnabás Szabó, Péter Marlok, István Péntek (National Food Chain Safety Office) and Dr Balázs Pataki for their help in the initial pedigree data collection and Andor Dallos† for his continuous help in collecting 16 years health data records in the stud.

